# Forecasting COVID-19 cases at the Amazon region: a comparison of classical and machine learning models

**DOI:** 10.1101/2020.10.09.332908

**Authors:** Dalton Garcia Borges de Souza, Francisco Tarcísio Alves Júnior, Nei Yoshihiro Soma

**Affiliations:** Collegiate of Industrial Engineering, University of Amapa State, Macapá, AP, Brazil; PROFNIT Program, PROFNIT-UNIFAP, Macapá, AP, Brazil; Division of Computer Science, Aeronautics Institute of Technology, São José dos Campos, SP, Brazil

## Abstract

**BACKGROUND:** Since the first reports of COVID-19, decision-makers have been using traditional epidemiological models to predict the days to come. However, the enhancement of computational power, the demand for adaptable predictive frameworks, the short past of the disease, and uncertainties related to input data and prediction rules, also make other classical and machine learning techniques viable options.

**OBJECTIVE:** This study investigates the efficiency of six models in forecasting COVID-19 confirmed cases with 17 days ahead. We compare the models autoregressive integrated moving average (ARIMA), Holt-Winters, support vector regression (SVR), k-nearest neighbors regressor (KNN), random trees regressor (RTR), seasonal linear regression with change-points (Prophet), and simple logistic regression (SLR).

**MATERIAL AND METHODS:** We implement the models to data provided by the health surveillance secretary of Amapáa, a Brazilian state fully carved in the Amazon rainforest, which has been experiencing high infection rates. We evaluate the models according to their capacity to forecast in different historical scenarios of the COVID-19 progression, such as exponential increases, sudden decreases, and stability periods of daily cases. To do so, we use a rolling forward splitting approach for out-of-sample validation. We employ the metrics RMSE, R-squared, and sMAPE in evaluating the model in different cross-validation sections.

**FINDINGS:** All models outperform SLG, especially Holt-Winters, that performs satisfactorily in all scenarios. SVR and ARIMA have better performances in isolated scenarios. To implement the comparisons, we have created a web application, which is available online.

**CONCLUSION:** This work represents an effort to assist the decision-makers of Amapá in future decisions to come, especially under scenarios of sudden variations in the number of confirmed cases of Amapá, which would be caused, for instance, by new contamination waves or vaccination. It is also an attempt to highlight alternative models that could be used in future epidemics.

## Introduction

By September 20^*th*^ 2020, almost nine months after SARS-COV-2 first appearance, World Health Organization (WHO) reported a total of 31.1 million cases worldwide, with 315, 919 daily cases and 962, 000 accumulated deaths due to coronavirus disease [1]. Brazilian authorities announced the first COVID-19 case by February 25^*th*^, 2020 [2]. Despite this two month delay, by the end of August 2020, Brazil already held the second largest number of accumulated infected (4.6 million) and death cases (137, 000) in the world. The number of daily new cases and deaths are also high, placing Brazil just after India and the United States of America, respectively, and both with much larger populations [3].

All those numbers caught the attention of many researchers, that presented models to attend the concerns from the Brazilian government and population, such as when the outbreak will peak, how long it will last, and how many will be infected or die [4, 5]. Many of those forecasting models rely on epidemiological approaches [6, 7] or state-of-art artificial intelligence (AI) algorithms. Generally, researchers address their models to the country as a unit or to highly populated areas, mainly big cities and federation states like Sao Paulo and Rio de Janeiro [4, 8, 9].

However, COVID-19 has also impacted other Brazilian regions, such as the North, that is a territory almost entirely covered by the Amazon rain-forest and accounts for almost half of the Brazilian territory. The north has a low population density (4.78 *inh*.*/km*^2^), accounts for only 8.8% of the Brazilian population, and is responsible for 14.3% of all confirmed cases of COVID-19 in Brazil. It may be represented by infected per population rates: 2.6% in the North, versus 1.5% in the rest of the country [10]. Figure 1 shows the evolution of the infection rate in all five Brazilian regions.

**Fig 1.**
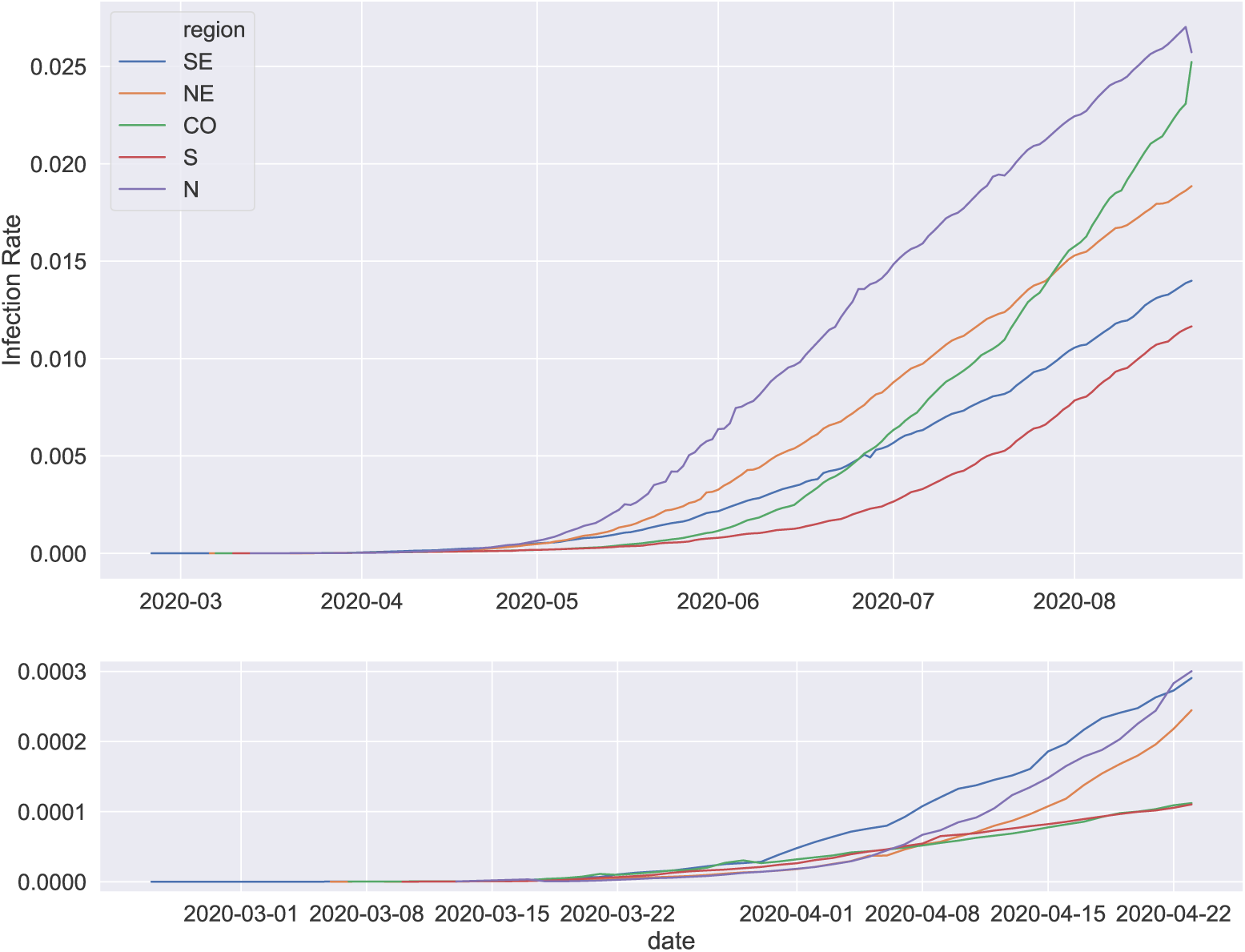
Evolution of the infection rate in all five Brazilian regions.

Carved into the Amazon rain-forest is Amapá, a northern state of Brazil. Amapá is like an island surrounded by the forest since it displays no land routes with any other Brazilian state (See Fig. 2). It has only 830,000 inhabitants but living in an area bigger than England, which is Voc67 times denser. Like other parts of Amazon, Amapá already experiences an excess mortality from infectious diseases, especially among indigenous populations. Despite recent political efforts, many people living in the state still suffers from different social and health problems such as minimal access to clean water and public sanitation [11]. Those and other reasons make Amapá especially susceptible to COVID-19 and other epidemic outbreaks that may occur in the future. By the end of May, Mapacá, the Amapá’s capital, saw its health system collapse due to COVID-19. By closing August 2020, the state consolidated the second highest infection rate in Brazil, according to official data [10]. By the end of September 2020, the state also has a low fatality rate (1.29%) when compared to the whole country (3.02%), which may be the result of local attempts to track new cases and avoid under-notifications.

**Fig 2.**
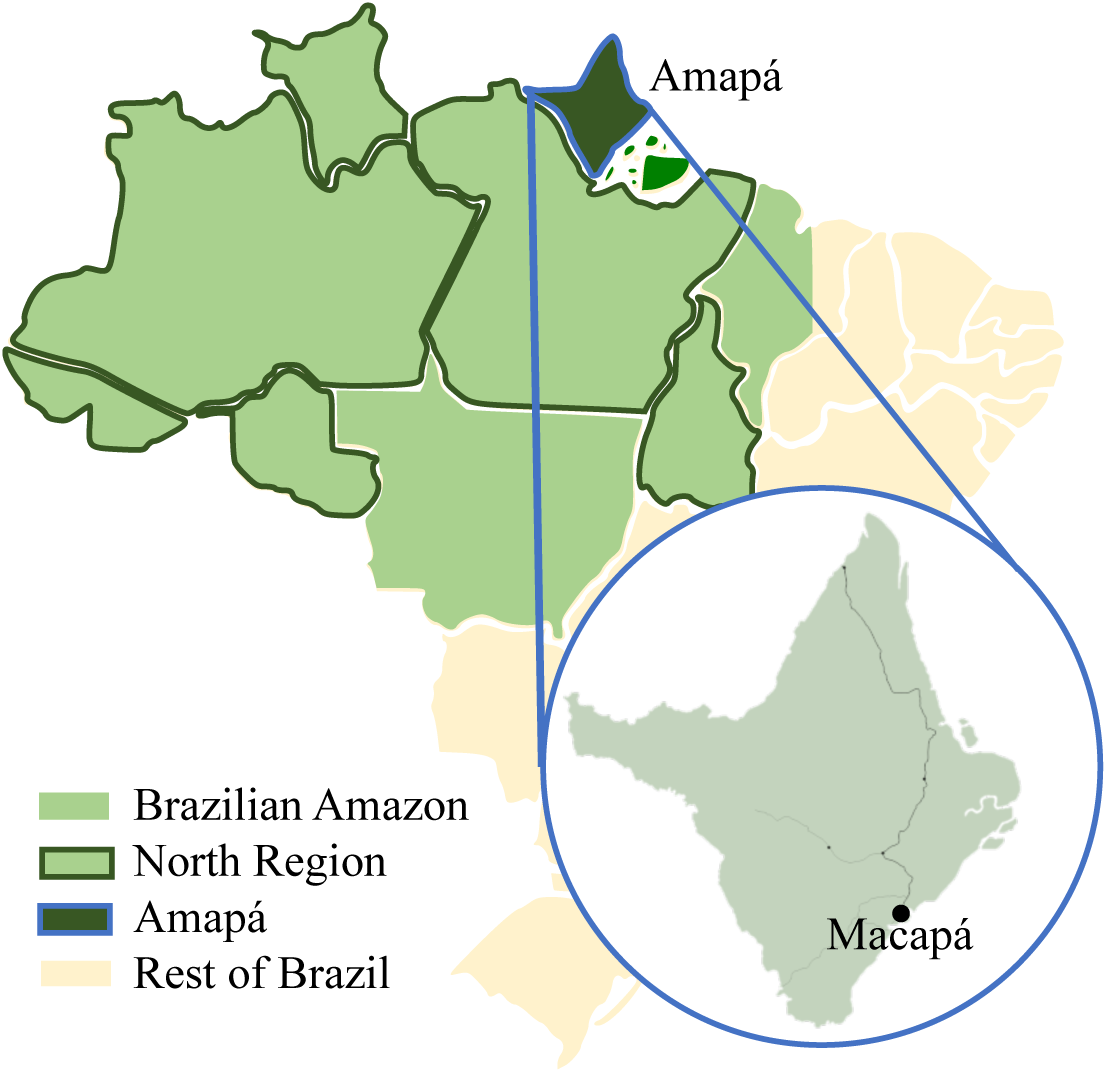
Amapá, Brazil.

Respecting this ambiance, in this paper, we explore and compare traditional and AI forecasting models to support the Amapaense decision-makers in the future decisions to come. The interest variables are the accumulated number of confirmed and death cases. We compare the models autoregressive integrated moving average (ARIMA), Holt-Winters, support vector regression (SVR), k-nearest neighbors regressor (KNN), random trees regressor (RT), seasonal linear regression with change-points (SLiR) and simple logistic regression (SLR), which dictates the baseline performance in this study. We compare the models according to the necessities of local authorities. Thus, we measure the model’s effectiveness to forecast the 17 days ahead and how fast they have responded to quick increases and decreases in the number of cases, as well as to periods of stability. This scenarios may repeat in the future, as result of new contamination waves or vaccination, for example. The forecasts are performed to each Amapaense municipality individually and to the state accumulated data, which we paint as our main example.

Since the municipalities are in different stages of the COVID-19 spreading, they may also display very different curve growing behaviors. Thus, as a result of this study, we have also created an online application (which can be accessed in http://www.previsor.covid19amapa.com/, that can be used to visualize the data at municipal level. The web application also allows decision-makers and researchers to follow the steps we do, as well as choose the best model to use in different occasions.

## 1 Research Framework

In this section, we describe our research framework, which we split into: (1.1) data acquisition, (1.2) data splitting, (1.3) fitting and forecasting, and (4) model evaluation. The subsection that follows treats each one of those steps.

### 1.1 Data acquisition

We performed all modelings to the cumulative confirmed cases of COVID-19 in Amapá, since the first official case, in March 20^*th*^ 2020, up to August 20^*th*^ 2020. We gather the data from official reports, from each of the 16^*th*^ Amapense municipalities. The collected data is also available in an application programming interface provided by Brasil.io repository [10]. The measurement periods are different for each municipality and Tab. 1 summarized the dates of the first and last reports.

**Table 1.** Number of observed days by city.

| Municipality | First report | Last report | Observ. days |
| --- | --- | --- | --- |
| Amapá | 26-Apr | 21-Sep | 149 |
| Calçoene | 1-May | 21-Sep | 144 |
| Cutias | 5-May | 21-Sep | 140 |
| Ferreira Gomes | 2-May | 21-Sep | 143 |
| Itaubal | 24-Apr | 21-Sep | 151 |
| Laranjal do Jari | 15-Apr | 21-Sep | 160 |
| Macapá | 20-Mar | 21-Sep | 186 |
| Mazegão | 14-Apr | 21-Sep | 161 |
| Oiapoque | 4-Apr | 21-Sep | 171 |
| Pedra Branca do Amapári | 23-Apr | 21-Sep | 152 |
| Porto Grande | 14-Apr | 21-Sep | 161 |
| Pracuúba | 5-May | 21-Sep | 140 |
| Santana | 5-Apr | 21-Sep | 170 |
| Serra do Navio | 22-Apr | 21-Sep | 153 |
| Tartarugalzinho | 26-Apr | 21-Sep | 149 |
| Vitória do Jari | 14-Apr | 21-Sep | 161 |

The data we use may diverge a little from the Brazilian government website, since the counting protocol may differ from those used by the Amapá state. Also, This paper does not treat case sub-notifications.

### 1.2 Data splitting

First, we split the raw data into training and test datasets. However, we performed a rolling forward splitting, with a minimum of *p* training samples and a fixed value of *q* testing samples. Considering a total of *n* observations, we first took *p* days as the training set and tried to forecast the next *q* days. Then, interactively, we added one day to the training set, until it comprised *n* − *q* observations. Thus, for a given municipality, we have *n* − *p* − *q* + 1 different cross-validation splittings.

In this paper, we use *p* and *q* equal to 17, since this the horizon required by Amapense decision-makers. Thus, in the first splitting, the raw data is divided into a proportion of half-and-half between training and testing sets (see Algorithm 1).

Each training sample (*x*) is then standardized (*z*) by its mean (*u*) and standard deviation (*s*), calculated as *z* = (*x* − *u*)*/s*.

#### Algorithm 1 cross-validation and forecasting

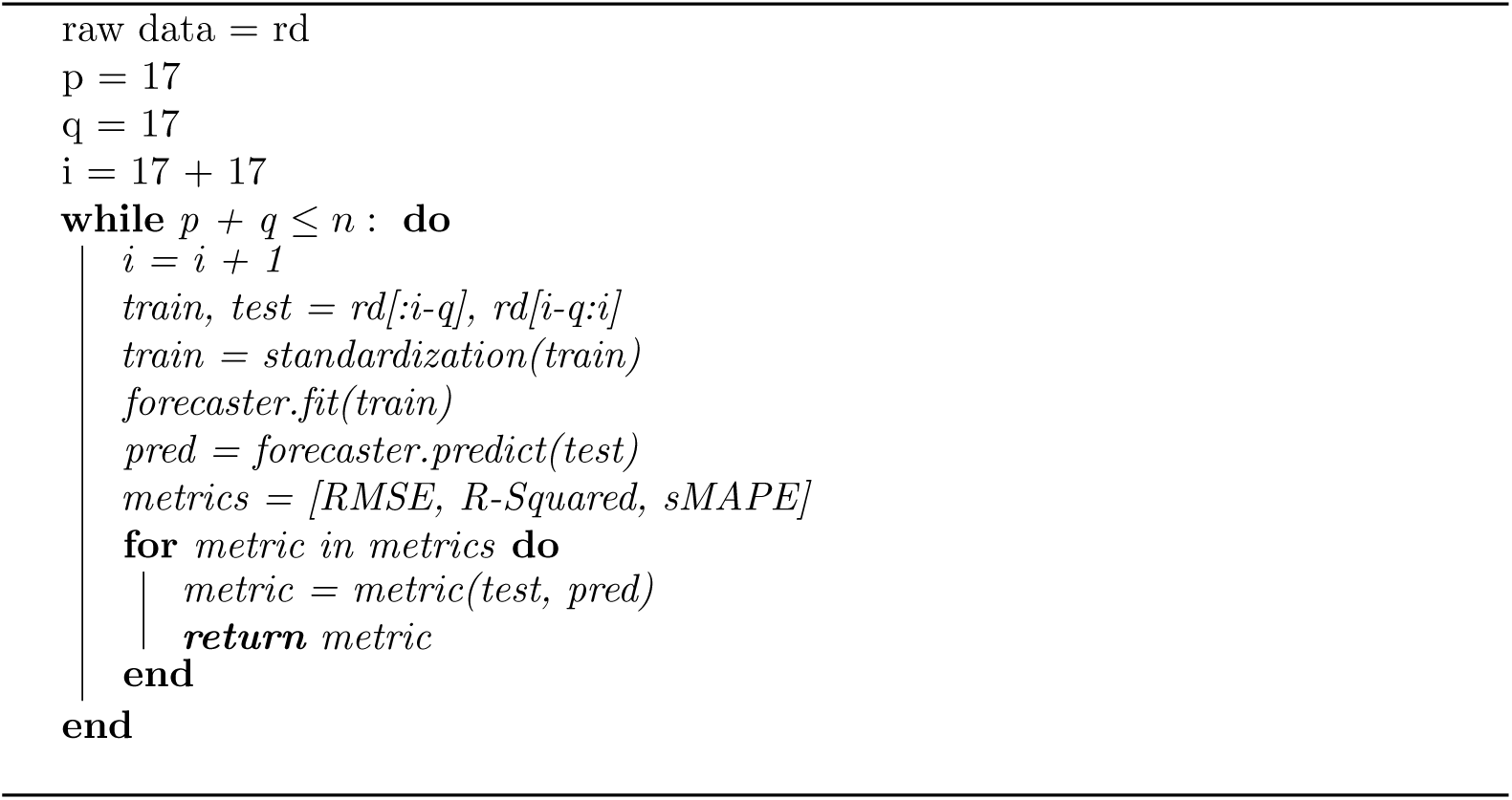

### 1.3 Fitting and Forecasting

We then fit the training datasets to each one the following models: autoregressive integrated moving average (ARIMA), Holt-Winters, support vector regression (SVR), k-nearest neighbors regressor (KNN), random forest regressor (RFR), seasonal linear regression with change-points (SLiR) and simple logistic regression (SLR). We also use search-grid to find the best hyperparameters sets to the state level data and for each model. With the exception of ARIMA model, which is automatized, when applicable the search-grid is performed only for the last time window we analyze. The models are applied to the bases that ensure the best fit for the model. Thus, the models Logistic Regression, Holt-Winters, ARIMA, and Prophet are modeled to accumulated databases, while SVR, KNN, and RFR use daily databases. we also reduce the Logistic, KNN, RFR and SVR regressors to the size of the testing samples, thus, 17 days. However, all of them are compared according to predicted accumulated values. This way, models running on a daily bases will convert the predicted values before calculating the metrics and comparing them. The models are explained as follows:

#### 1.3.1 ARIMA

The ARIMA model stands for integration (I) between autoregressive (AR) and moving average (MA) models. Box and Jerkings [12] are the first designers of this model. ARIMA may also be adjusted to consider seasonality, which optimal value may be found after the conduction of a Canova-Hansen test [13]. The optimum values of autoregressive (*p*), degree of their differences (*d*) and moving average (*q*) may also be found by search-grid. Usually, we select the parameters that minimize the Information Criterion (AIC). Articles such as Benvenuto et al. [14], Ceylan [15], and Singh et al. [16] bring examples of ARIMA applications to COVID-19 cases forecasting. The general equations for AR and MA models are [15]:

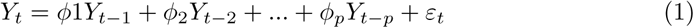

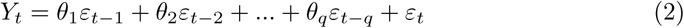

where *Y*_*t*_, *ε, φ*, and *θ* are the observed values at time *t*, the value of the random shock at time *t*, AR, and MA parameters, respectively. Thus, an ARMA model is given by:

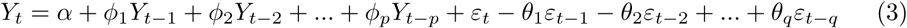

Where *α* is a constant. When dealing with non stationarity, the data may be differenced, and the ARIMA model is then performed.

#### 1.3.2 Holt-Winters

Holt [17] and Winters [18] are the architectures of the Holt-Winters method, also known as triple exponential smoothing. This model is an upgraded version of the simple exponential smoothing to consider trend and seasonality. Thus, it employs three parameters: *α*, the smoothing factor, *β*, a trend smoothing parameter, and *γ*, which relates to seasonality. Different authors have explored this model to forecast COVID-19 cases [19, 20]. The equations of the additive model follow.

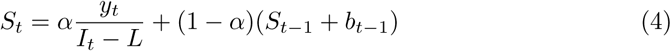

Where *S* is the smoothed observation, *L* the cycle length, and *t* a period. The trend factor (*b*), the seasonal index (*I*), and the forecast at *m* steps (*F*) are given by:

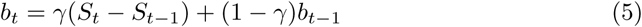

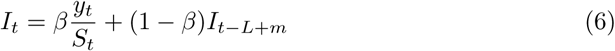

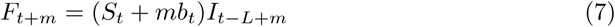

#### 1.3.3 SVR

A support vector machine (SVM) is a supervised machine learning algorithm. It performs both regression and classification tasks. Vapnik [21] is the precursor of this technique, and its variant for regression, the support vector regression (SVR), which was widespread mainly by the work of Drucker et al. [22]. Some applications of SVR can be found in the context of COVID-19 case forecasting [4, 23, 24].

The general logic of an SVR is relatively simple. Suppose a linear regression, which objective is to minimize the sum of square errors.

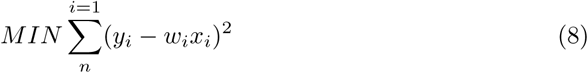

where *y*_*i*_ is the target, *w*_*i*_ the coefficient and *x*_*i*_ the feature. Then, the training of SVR aims to minimize the following system.

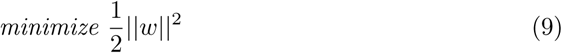

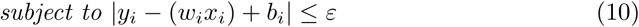

#### 1.3.4 KNN

KNN stands for k-nearest neighbors and was primarily designed to deal with classification problems. Decades after the first conceptualizations of KNN, around the start of the’90s, researchers started exploring it for regression purposes [25]. In the time series context, the KNN algorithm searches for k nearest past similar values by minimizing a similarity measure. Then, the forecasting is an average of these k-nearest neighbors. However, it sounds straightforward, it demands a high computational cost [9]. In the context of COVID-19, Many researchers have used this approach in classification problems. Just a few have used it to COVID-19 case forecasting [9]. The main distance functions used for continuous variables are:

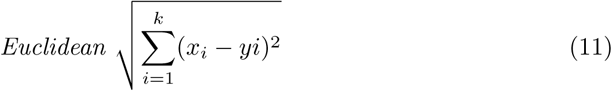

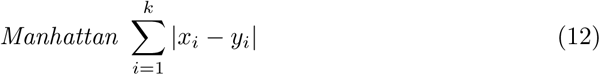

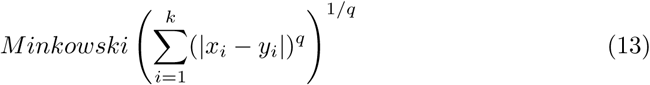

#### 1.3.5 RFR

Random forest is a machine learning algorithm with many decision trees. Breiman [26] proposed a combination of bagging and random subspaces methods. Nowadays, researchers and machine learning practitioners employ RF in both classification and regression tasks. Authors have applied RF to deal with COVID-19 forecasting [4, 27]. This model randomly splits the data into in-Bag data and out-of-Bag data. Then many decision trees are randomly created with bootstrap samples. The branching of each tree is also performed according to randomly selected predictors. The final RF estimate in an average of the results from each tree. It is especially impressive when dealing with the randomness of the time series. In regression applications, Mean Squared Error (MSE) is used as splitting criteria in each tree’s branch. We explain MSE in more detail later on.

#### 1.3.6 Prophet

Prophet is a forecasting approach developed by Facebook. It employs a decomposable times series model, with three main model components: trend (*g*(*t*)), seasonality (*s*(*t*)) and holidays (*h*(*t*)). It also assumes an error *ϵ* representing any idiosyncratic changes that are not predicted by the model.

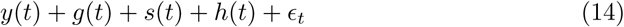

with

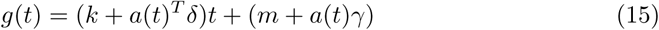

where *k* is the growth rate, *δ* is the rate adjustments, *m* is the offset parameter, and *γ*_*j*_ is set to *s*_*j*_*δ*_*j*_ to make the function continuous. Another important aspect is that the model performs automatic changepoint selection, putting a sparse prior on *δ*.

On the other hand, it relies on Fourier series to incorporate daily, weekly, and annually seasonalities. In the case of COVID-19, we are more concerned about weekly seasonality.

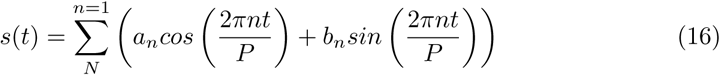

In the context of COVID-19, Prophet has few appearances in forecasting the accumulated confirmed and death cases [8, 28].

### 1.4 Model evaluation

We evaluate the performance of each forecasting models in terms of R-squared (*R*^2^), Root Mean Square Error (RMSE), and Symmetric Mean Absolute Percentage Error (SMAPE). We perform the evaluations for each train/test pair created by the rolling forward splitting. Thus, each metric is performed *n* − *p* − *q* + 1 times.

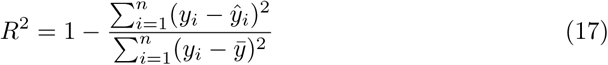

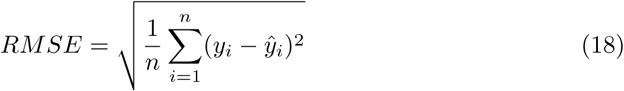

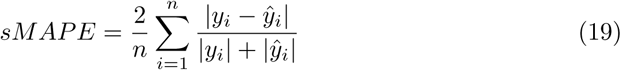

where n is the number of observations, *y*_*i*_ and *ŷ*_*i*_ are the *i*^*th*^ observed and predicted values.

## 2 Results and Discussion

This section describes the results of our experiments. We compare the models according to their efficiency to predict COVID-19 confirmed cases 17 days ahead and over different time windows. In the next subsections, we describe the models’ performances during exponential increase, after a sudden decrease, during the stability of daily new cases, and overall for the whole period.

### 2.1 Exponential increase

For the data related to the State as a whole, we take the period between March 20^*th*^ and June 20^*th*^ as an example of period with exponential increase. Among all models, ARIMA is the one that seems to perform better in the period, considering 17 days of rolling forward windows from May 1^*st*^ to June 20*th*. Holt-Winters follows ARIMA during this period and even displaying slightly better results in the last 17 days window (ARIMA: RMSE = 649, R-Squared = 0.95, and sMAPE = 3.81; Holt-Winters: RMSE = 575, R-Squared = 0.96, and sMAPE = 3.09). Fig. 3 shows how Holt-Winters graphically fits the data.

**Fig 3.**
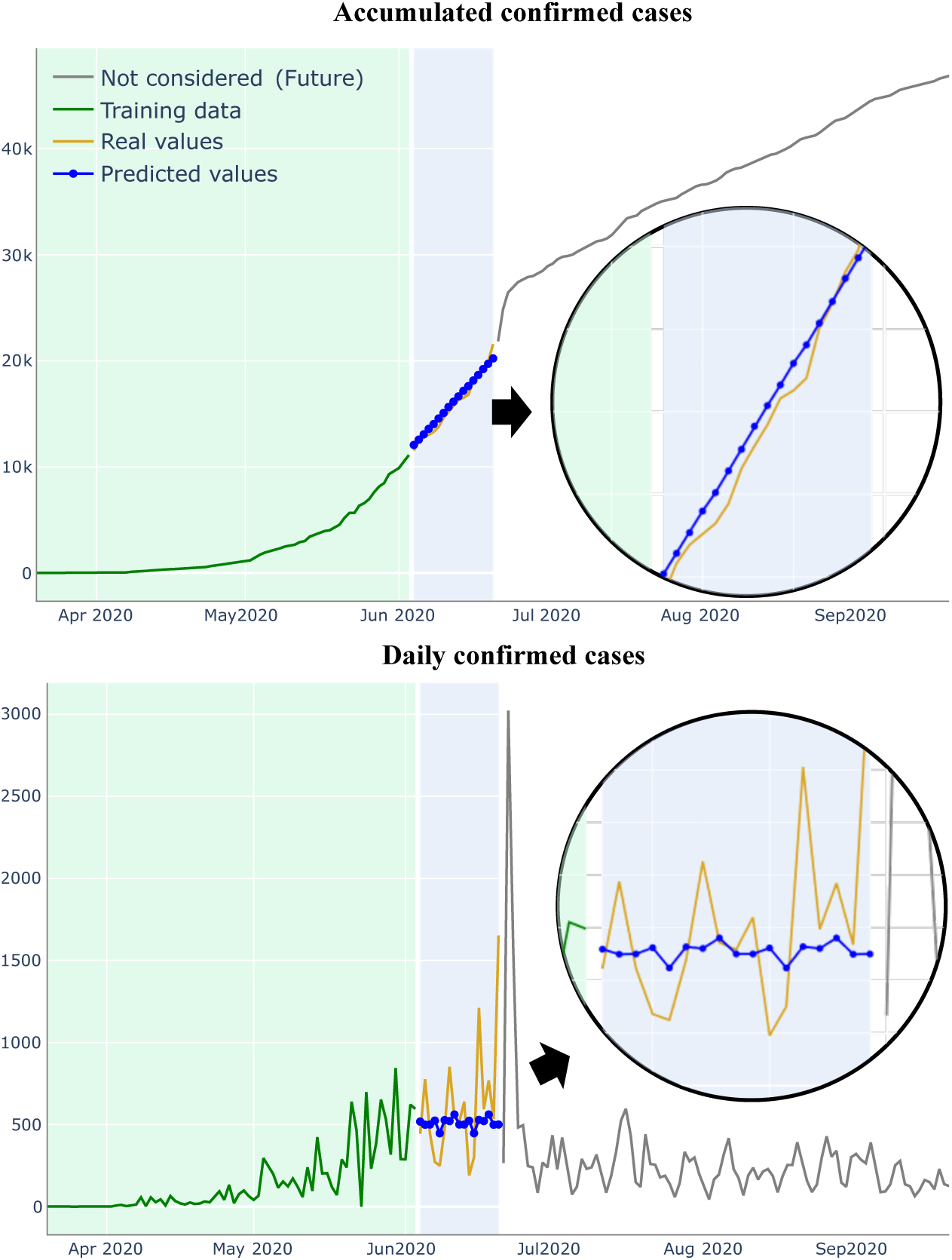
Forecasting during exponential growth with Holt-Winters.

### 2.2 After sudden decrease

Still, for the Amapá state, we take the period between June 22*th* and July 14^*th*^ as an example of a period after a sudden decrease. In this case, Holt-Winters, ARIMA, and RFR are those that perform better, considering 17 days rolling forward windows. For instance, in the last 17 days window Holt-Winters displays RMSE = 262, R-Squared = 0.95, and sMAPE = 0.74. Fig. 4 shows how Holt-Winters graphically fits this kind of data. Holt-Winters has a good performance for this specific time window. All forecasting models struggle to predict just after a sudden decrease in the number of daily confirmed cases. The models that have the fastest recover are SVR and Holt-Winters, in this order.

**Fig 4.**
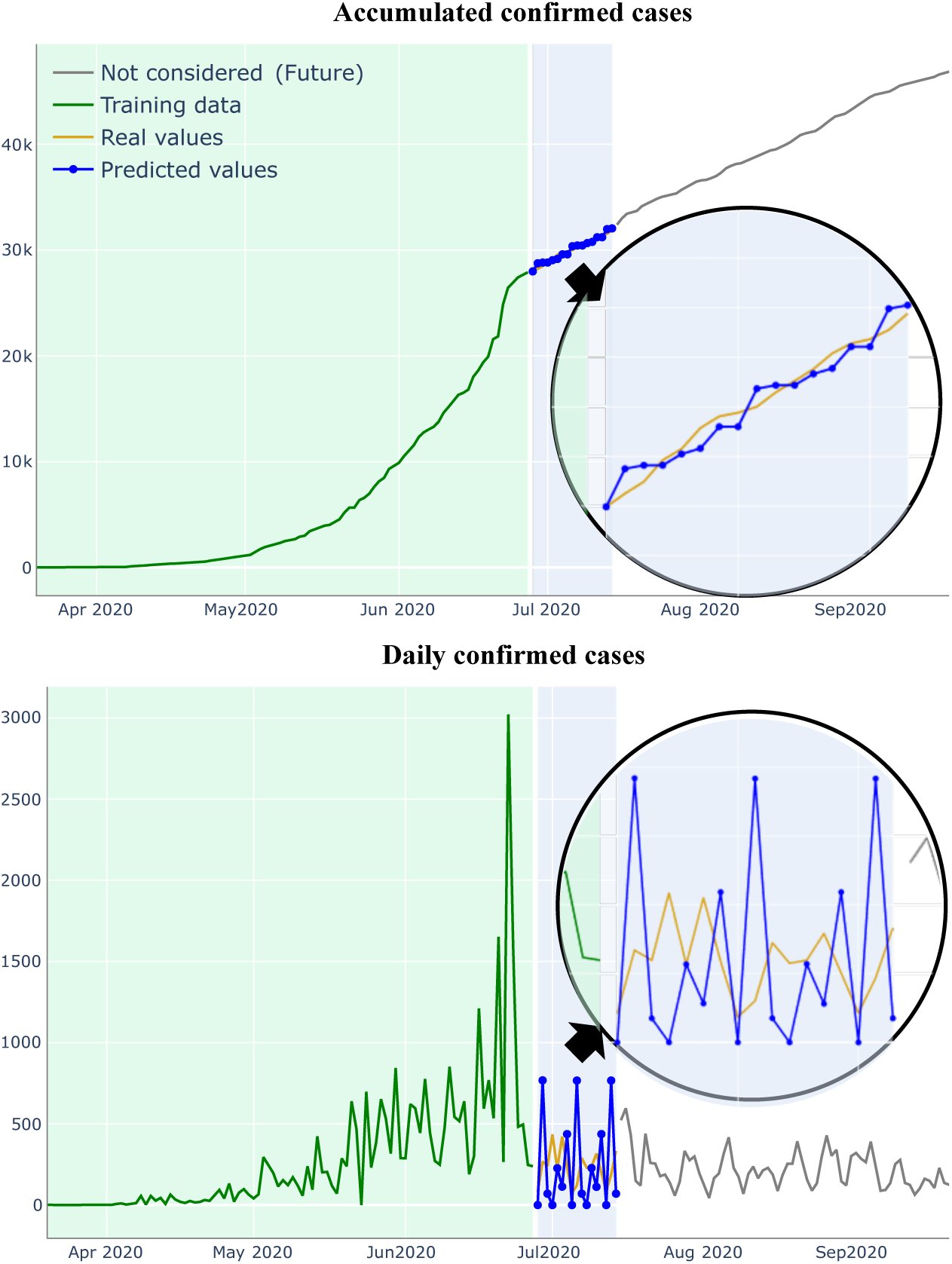
Forecasting during stability with Holt-Winters.

### 2.3 Stability period

We take the period between July 14^*th*^ to September 20^*th*^ as a period of stability in daily new cases, where the average of daily new cases tends to be constant, and weekly seasonal variations and noise mostly influence the values. In this case, all ML models perform reasonably, with special attention to Holt-Winters, ARIMA, and SVR. For instance, Fig. 5 shows hot Holt-Winters graphically fits to a section of this type of period (RMSE = 162, R-Squared = 0.98, sMAPE = 0.34). At the same section, Logistic regression display as metrics RMSE = 667, R-Squared = 0.6 and sMAPE = 1.34.

**Fig 5.**
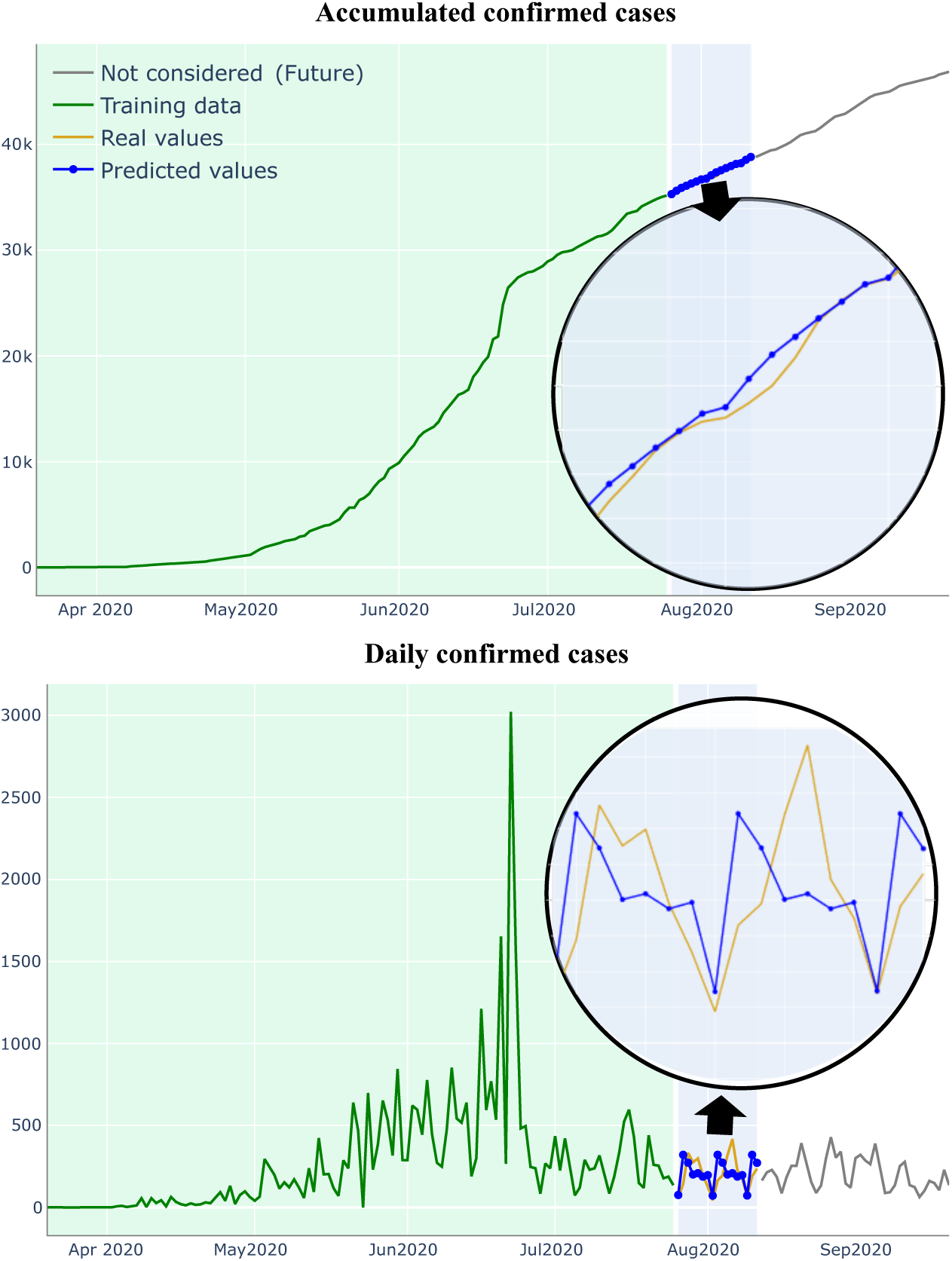
Forecasting during stability with Holt-Winters.

### 2.4 Model’s overall performance

In a general manner, all machine Learning models achieve better results than Logistic regression. In Fig. 6 we can see how Holt-Winters performs in comparison to the other five models. Those findings Notice that we measure rolling forward performances according to the R-Squared given by each cross-validation set.

**Fig 6.**
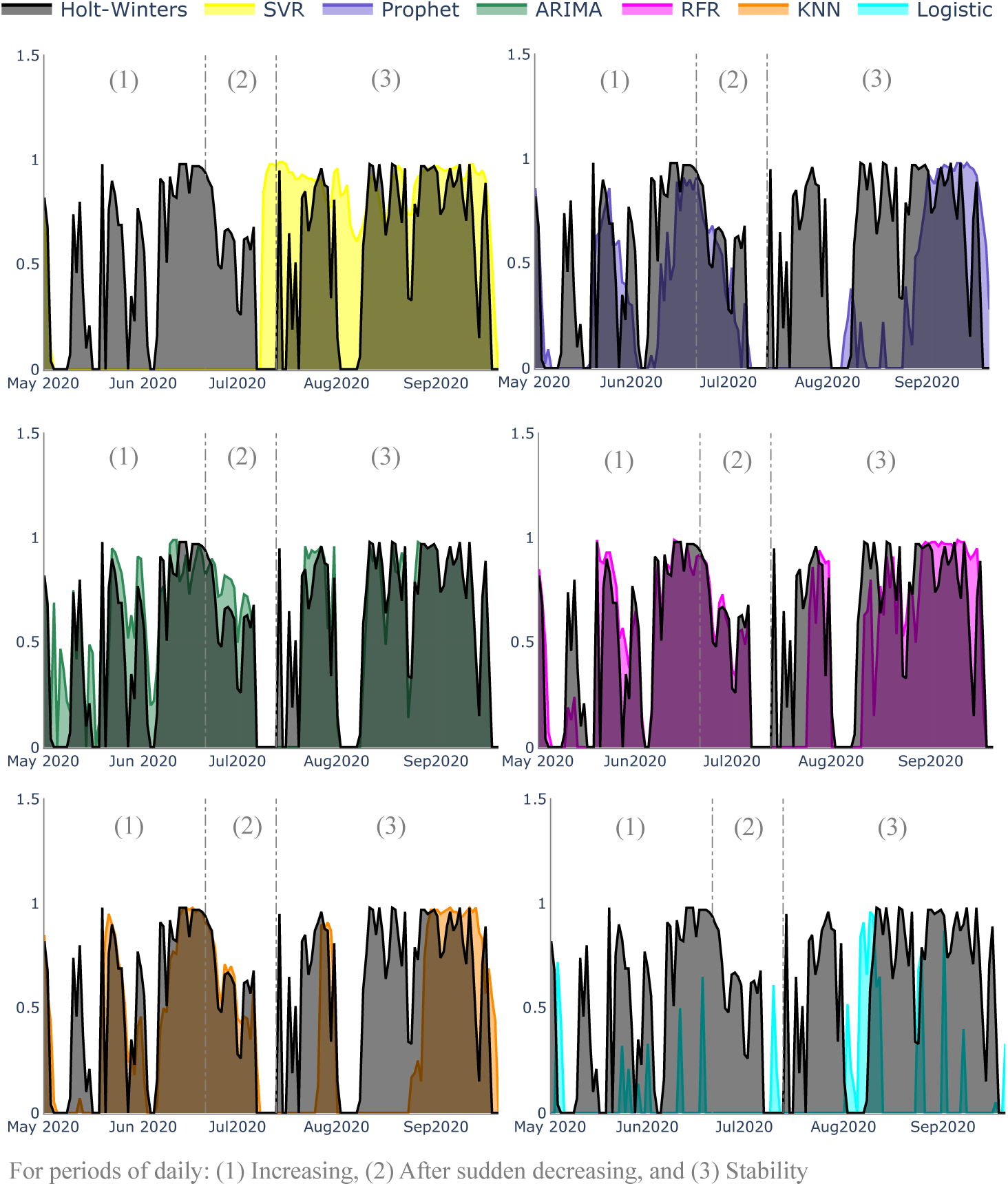
Rolling forward performance of Holt-Winters in comparison to other models (R-squared metric)

In each pair of models we can observe how the Holt-Winters perform in comparison to an other model and considering the periods we classify as (1) exponential increasing, (2) after sudden daily decreasing and (3) stability of daily new cases.

Similar evaluations to the prediction of confirmed cases can be extended to death cases. In this case, Holt-Winters still seems to be the most suitable model, along with ARIMA. Similar considerations can also be draw to the municipalities of Amapá. However, for small cities where data is scarce, most models we present here struggle to make predictions. In this case, even naive approaches seem to be a good alternative.

## 3 Conclusions

COVID-19 has been a burden issue to the world, and to a large number of countries. It imposes severe challenges to local authorities that do not have the necessary resources to fight it. The situation of Amapá is not different, a Brazilian state in the middle of the Amazon rain-forest. Like other Amazonian regions, various Amapaense communities already suffer from social and health problems, such as minimal access to public sanitation and different epidemiological diseases, such as Malaria and Yellow Fever. COVID-19 depreciates these social conditions. Knowing how the COVID-19 numbers will evolve is critical to local authorities to determine the best responses.

Thus, in this paper, we compared classical and machine learning models to forecast the evolution of COVID-19 in the state. Despite the volume of research papers pointing Machine Learning models as those with the best performance for many locations, in the case of Amapá, two classical approaches seem to perform better: Holt-Winters and ARIMA. It may be a consequence of the Amapaense data, which has marked seasonality and sudden variations. One advantage of these two models is that they are easier to code and tune than machine learning models.

This conclusions, as well as other analysis, can be made by exploring the web application we created and is available online at http://www.previsor.covid19amapa.com/.

As possible developments of this research, we highlight the investigation of Neural Networks models, which may consider other feature sets in forecasting future numbers of cases. We also intend to propose a framework that indicates the best forecasting model for each municipality and period, saving time from local decision-makers.

## 4 Acknowledgments

The authors would like to thank the Universidade Estadual do Amapá (UEAP), for the financial and administrative structure provided.

